# ZIKV disrupts placental ultrastructure and drug transporter expression in mice

**DOI:** 10.1101/2020.12.17.423218

**Authors:** Cherley Borba Vieira de Andrade, Victoria Regina de Siqueira Monteiro, Sharton Vinicius Antunes Coelho, Hanailly Ribeiro Gomes, Ronny Paiva Campos de Sousa, Veronica Muller de Oliveira Nascimento, Flavia Fonseca Bloise, Stephen Matthews, Enrrico Bloise, Luciana Barros de Arruda, Tania Maria Ortiga-Carvalho

## Abstract

Congenital Zika virus (ZIKV) infection can induce fetal brain abnormalities. Here, we investigated whether maternal ZIKV infection affects placental physiology and metabolic transport potential and impacts the fetal outcome, regardless of viral presence in the fetus at term. Low (10^3^ PFU-ZIKVPE243; low ZIKV) and high (5×10^7^ PFU-ZIKVPE243; high ZIKV) virus titers were injected into immunocompetent (ICompetent C57BL/6) and immunocompromised (ICompromised A129) mice at gestational day (GD) 12.5 for tissue collection at GD18.5 (term). High ZIKV elicited fetal death rates of 66% and 100%, whereas low ZIKV induced fetal death rates of 0% and 60% in C57BL/6 and A129 dams, respectively. All surviving fetuses exhibited intrauterine growth restriction (IUGR) and decreased placental efficiency. High-ZIKV infection in C57BL/6 and A129 mice resulted in virus detection in maternal spleens and placenta, but only A129 fetuses presented virus RNA in the brain. Nevertheless, pregnancies in both strains produced fetuses with decreased head sizes (p<0.05). Low-ZIKV-A129 dams had higher IL-6 and CXCL1 levels (p<0.05), and their placentas showed increased CCL-2 and CXCL-1 contents (p<0.05). In contrast, low-ZIKV-C57BL/6 dams had an elevated CCL2 serum level and increased type I and II IFN expression in the placenta. Notably, less abundant microvilli and mitochondrial degeneration were evidenced in the placental labyrinth zone (Lz) of ICompromised and high-ZIKV-ICompetent mice but not in low-ZIKV-C57BL/6 mice. In addition, decreased placental expression of the drug transporters P-glycoprotein (P-gp) and breast cancer resistance protein (Bcrp) and the lipid transporter Abca1 was detected in all ZIKV-infected groups, but Bcrp and Abca1 were only reduced in ICompromised and high-ZIKV ICompetent mice. Our data indicate that gestational ZIKV infection triggers specific proinflammatory responses and affects placental turnover and transporter expression in a manner dependent on virus concentration and maternal immune status. Placental damage may impair proper fetal-maternal exchange function and fetal growth/survival, likely contributing to congenital Zika syndrome.

## 1. Introduction

Congenital Zika virus (ZIKV) infection can be associated with adverse pregnancy outcomes. Neonates born from ZIKV-positive pregnancies may develop severe neurological abnormalities, placental pathologies and intrauterine growth restriction (IUGR), among other complications (1). ZIKV vertical transmission has become a major public health issue worldwide, especially in Brazil, where more than 200,000 ZIKV-positive cases have been confirmed and over 2,000 congenital microcephaly births have been reported (2–6). These numbers represent a 20-fold rise in the incidence of congenital microcephaly in Brazil during the years of the ZIKV pandemic, with similar increases reported elsewhere in Latin America (2,3,7). Importantly, while the ZIKV pandemic is currently thought to be controlled, evidence points to a possible silent ZIKV spread across the Americas (8,9), highlighting the need for improved knowledge of the possible routes of vertical ZIKV transmission and its association with disruptive inflammatory and developmental phenotypes and the need for new avenues of prevention and treatment.

Previous studies have investigated the possible pathways involved in vertical ZIKV transmission. Miranda and colleagues (10) showed that in humans, ZIKV infection changed the pattern of tight junction proteins, such as claudin-4, in syncytiotrophoblasts. Jurado et al. (2016) suggested that the migratory activities of Hofbauer cells (feto-placental macrophages) could help disseminate ZIKV to the fetal brain (11). Other recent studies have shown that placental villous fibroblasts, cytotrophoblasts, endothelial cells and Hofbauer cells are permissive to ZIKV, and placentae from ZIKV-infected women had chorionic villi with a high mean diameter (11–14). Furthermore, in 2019, Rathore et al. demonstrated that pregnant mice carrying high levels of antibodies against dengue virus (DENV) exhibited increased ZIKV vertical transmission associated with severe microcephaly-like syndrome, demonstrating another possible mechanism of antibody-dependent vertical ZIKV transmission (15). However, at present, further studies are required to identify the precise mechanism of maternal-fetal ZIKV transmission.

Many mouse models have been developed to identify how ZIKV overcomes placental defenses. Initially, limited information was obtained due to the apparent inability of the virus to infect wild-type (WT) mice (16). ZIKV NS5 targets the interferon signaling pathway in humans but not in mice (17). Thus, WT mice show no clear evidence of clinical disease (17,18) and are of limited use in modeling the disease. However, mice lacking an interferon signaling response show evidence of disease and have been widely used to investigate ZIKV infection during pregnancy (8,17,19).

The interferon system, especially type III interferon, is a key mechanism of host defense and a viral target for immune evasion (20). Type III interferons have a role in protection against ZIKV infection in human syncytiotrophoblasts from term placenta (21). Luo et al. have shown that inhibition of Toll-like receptors 3 and 8 inhibits the cytokine output of ZIKV-infected trophoblasts (22). In addition, viral replication coincides with the induction of proinflammatory cytokines, such as interleukin [IL]-6. This cytokine has a crucial role in inflammation and affects the homeostatic processes related to tissue injury and activation of stress-related responses (23,24). ZIKV infection can trigger an inflammatory response with IL-6 release (11,25).

Maternal infection has profound effects on placental permeability to drugs and environmental toxins. Changes in the expression and function of specific ABC transporters in the placenta and yolk sac following infective and inflammatory stimuli have been demonstrated (26–30). ABC transporters are efflux transporters that control the biodistribution of several endogenous and exogenous substrates, including xenobiotics (antiretrovirals and synthetic glucocorticoids), steroid hormones (estrogens and androgens), nutrients (folate and cholesterol) and immunological factors (chemokines and cytokines) within the maternal-fetal interface (31). The best described ABC transporters in the placenta are P-glycoprotein (P-gp; also known as multidrug resistance protein 1, MDR1), breast cancer resistance protein (Bcrp) and the lipid Abca1 and Abcg1 transporters. P-gp and Bcrp transporters are responsible for preventing fetal accumulation of xenobiotics and environmental toxins that may be present in the maternal circulation, whereas Abca1 and Abcg1 control the placental exchange of cytotoxic oxysterol and lipid permeability throughout pregnancy; therefore, they play an important role in fetal protection and placental lipid homeostasis (26).

Despite the limited number of studies showing ZIKV infection in immunocompetent mice, intrauterine inoculation with a high virus titer was previously demonstrated to result in decreased fetal viability, with worse outcomes following infection in early gestation (32). In another report, intravenous infection on a very early embryonic day resulted in fetal demise even though the virus was not found in the fetal compartment in most of the treated animals (33). In the present study, we hypothesize that maternal exposure to ZIKV affects placental function, including placental ultrastructure and ABC transporter (P-gp, Bcrp, Abca1 and Abcg1) protein expression, even in the absence of vertical transmission and that these effects are dependent on viral infective titers and maternal immune status.

## 2. Materials and methods

### 2.1 Virus preparation and storage

The Brazilian ZIKV_PE243_ (GenBank ref. number KX197192) strain was isolated from a febrile case during the ZIKV outbreak in the state of Pernambuco, Brazil and was kindly provided by Dr Ernesto T. Marques Jr. (Centro de Pesquisa Aggeu Magalhães, FIOCRUZ, PE). Viruses were propagated in C6/36 cells, and viral titers were determined by plaque assays in Vero cells, as previously described (34). Supernatants of noninfected C6/36 cells cultured under the same conditions were used as mock controls.

### 2.2 Animal experimentation and study design

Two mouse strains were used in the study: immunocompetent (ICompetent) C57BL/6 and immunocompromised (ICompromised) (type 1 *Ifnr*-deficient) A129 strains. Since we were unable to consistently produce viable pregnancies by mating A129 males and females in our experimental settings, we mated A129 females (n=15) with C57BL/6 males (n=4) to produce ICompromised C57BL6/A129 pregnancies, whereas ICompetent C57BL/6 pregnancies were obtained by mating male (n=6) and female (n=35) C57BL/6 mice (8-10 weeks old). Animals were kept in a controlled temperature room (23°C) with a light/dark cycle of 12 hours and *ad libitum* access to water and food. After detection of the proestrous/estrous phase via vaginal cytology, copulation was confirmed by visualization of the vaginal plug and considered gestational day 0.5 (GD0.5). Maternal weight was monitored for confirmation of pregnancy; thus, females were weighed on GD0.5 and GD12.5, and females with a weight gain greater than 3 g were considered pregnant and entered randomly in the experimental groups. Experimental protocols were approved by the Animal Care Committee of the Health Sciences Center, Federal University of Rio de Janeiro (CEUA-036/16 and 104/16) and registered with the Brazilian National Council for Animal Experimentation Control.

On GD12.5, pregnant mice (ICompetent and ICompromised pregnancies) were injected with a single intravenous (i.v.) titer of ZIKV or mock control. ICompetent and ICompromised pregnant mice were randomly subdivided into three experimental groups: the mock (control) group, which received an injection of supernatant from noninfected C6/36 cells (ICompetent mock and ICompromised mock); the high-ZIKV-titer group, inoculated with 5×10^7^ plaque-forming units (PFU) of ZIKV_PE243_ (ICompetent high and ICompromised high); and the ZIKV low-titer group, injected with 10^3^ PFU of ZIKV_PE243_ (ICompetent low and ICompromised low).

On the morning of GD18.5, all animals were euthanized with a sodium phenobarbital overdose of 300 mg/kg. Maternal blood was collected via cardiac puncture, centrifuged (10 min, 4000 g) and stored at −20°C. The maternal brain and spleen and all placentae and all fetuses were dissected, collected and weighed, followed by fetal head isolation and measurement. The three placentae closest to the mean weight in a litter were selected for further analysis and cut in half using umbilical cord insertion as a reference (35–37). One-half of the placental disk was frozen in liquid nitrogen for qPCR, and the other half was fixed overnight in buffered paraformaldehyde (4%, Sigma-Aldrich, Brazil) for ultrastructural and protein expression/localization analysis. Matched fetal heads, maternal brains and spleens were frozen in liquid nitrogen for qPCR. Of important note, all fetuses obtained from ICompromised pregnancies were heterozygous.

### 2.3 ZIKV RAN quantification via RT-qPCR

ZIKV load was evaluated in maternal blood, brains and spleens and in the placentae and fetal heads. Brains, spleens, placentae and fetal heads were macerated in RPMI medium (Gibco™ RPMI 1640 Medium) normalized by the ratio of 0.2 mg of tissue to every 1 μl of medium and plotted per gram of tissue. The macerated volume was centrifuged at 4500 g for 5 min to remove tissue residues, and then, 500 μl of the centrifuged volume was used for RNA extraction using 1 mL of TRIzol reagent (Life Technologies, Thermo Fischer, USA). Treatment with DNase I (Ambion, Thermo Fischer, USA) was performed to prevent contamination by genomic DNA. cDNA was synthesized using a cDNA High Capacity Kit (Applied Biosystems, Thermo Fischer, USA) according to the manufacturer’s instructions by subjecting the samples to the following cycle: 25°C for 10 min, 37°C for 120 min and 85°C for 5 min. qPCR was performed using a StepOnePlus Real-Time qPCR system, TaqMan Master Mix Reagents (Applied Biosystems, Thermo Fischer, USA) and primers and probes specific for the protein E sequence (38). Samples were then subjected to the following cycle: 50°C for 2 min, followed by 40 cycles of 95°C for 10 min, 95°C for 15 sec, and 60°C for 1 min.

#### 2.3.1 RT-qPCR

The placenta was macerated in 1.5 mL of TRIzol reagent (Life Technologies, Thermo Fischer, USA). RNA extraction was performed following the manufacturer’s protocol. cDNA was prepared using Power SYBR Green PCR Master Mix (Life Technologies, Thermo Fisher, USA). The reaction was carried out for selected genes using intron-spanning primers (Table 1) and the StepOnePlus Real-Time PCR system (Life Technologies, Thermo Fischer, USA). Samples were subjected to the following cycle: 95°C for 10 min, followed by 40 amplification cycles consisting of DNA denaturation for 30 sec at 95°C and annealing of primers for 30 sec at 60°C. The threshold cycle (Ct) was determined for each gene of interest and for the reference genes glycerol 3-phosphate dehydrogenase (*Gapdh*) and RNA Polymerase II Subunit A (*Polr2a*). The relative expression of each gene was calculated using 2^−ΔΔCT^ (39) and graphically expressed as the fold-increase. The efficiency was calculated using the standard curve method. The melting curves were analyzed for each sample.

**Table 1:**
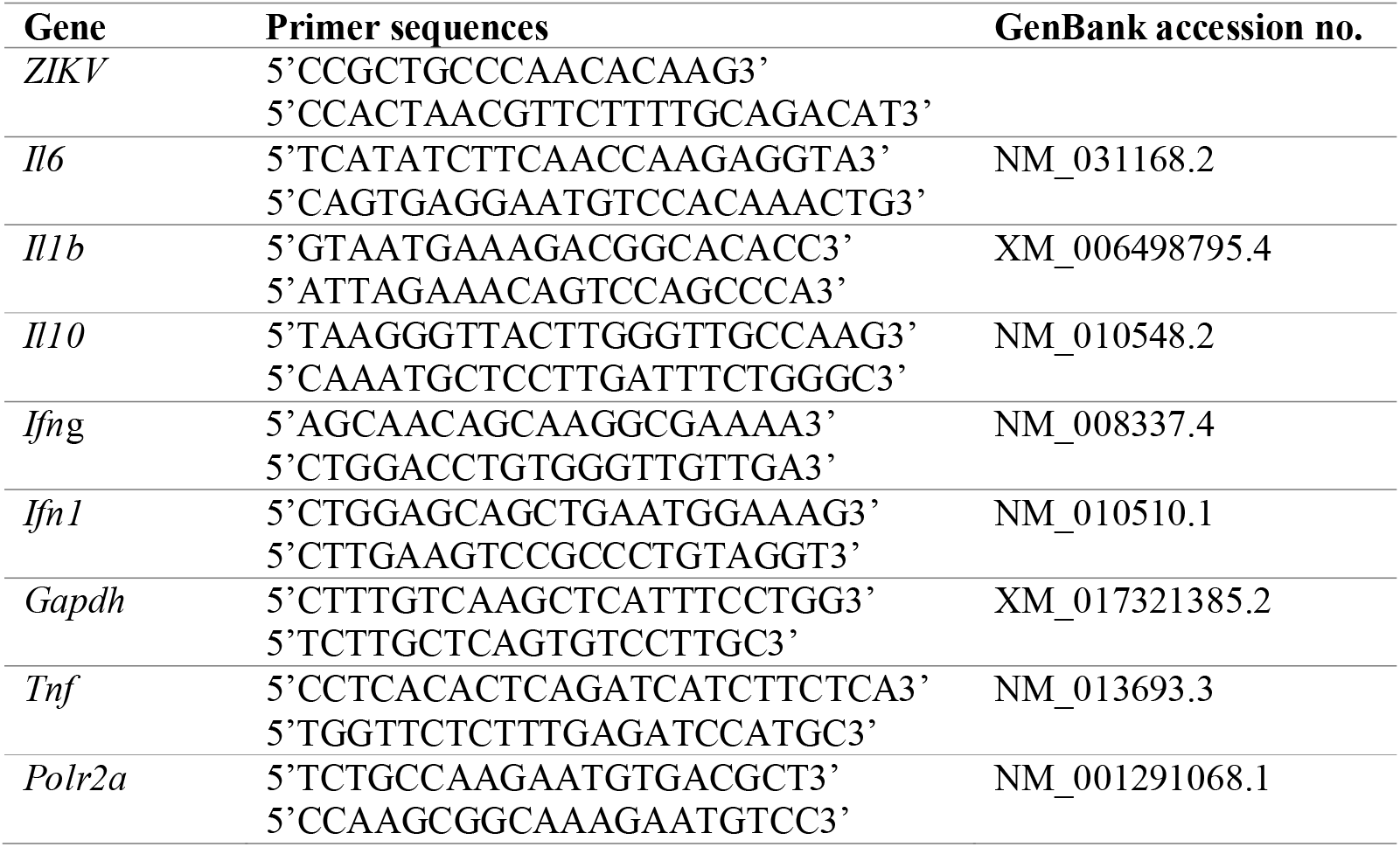
Primer sequences for the real-time PCR assay.

### 2.4 Detection of cytokines and chemokines in maternal serum and the placenta

Initially, placental tissue was homogenized in extraction buffer (50 mM Tris, 150 mM NaCl, 1X Triton, 0.1% SDS, 5 mM EDTA, 5 mM NaF, 50 mM sodium pyrophosphate, 1 mM sodium orthovanadate, pH 7.4) containing complete protease inhibitor cocktail (Roche Applied Science, Germany) with TissueLyser LT (Qiagen, Germany). The protein concentration of each sample was analyzed using a Pierce™ BCA Protein Assay Kit (Thermo Scientific, USA) according to the manufacturer’s instructions. Analysis of the cytokines IL-6 and IL-1β and the chemokines monocyte chemoattractant protein-1 (MCP-1/CCL2) and chemokine (C-X-C motif) ligand 1 (CXCL1) in maternal serum and placenta was performed with MILLIPLEX MAP Mouse Cytokine/Chemokine Magnetic Bead Panel – Immunology Multiplex Assays (MCYTOMAG-70K, Merck Millipore, Germany) following the manufacturer’s recommendations. The plate with samples and magnetic beads was analyzed on a MAGPIX^®^ System (Merck Millipore, Germany). The analyses were performed using Luminex xPonent^®^□ for MAGPIX^®^□ v software. 4.2 (Luminex Corp., USA). For each reaction well, the MAGPIX Luminex^®^ platform reports the median fluorescence intensity (MFI) for each of the analytes in the sample. The levels of each analyte were then calculated against the standard curve. The ratio between the value obtained and the protein quantification for each sample was determined and plotted.

### 2.5 Virus titration using plaque assays

Blood from mock- and ZIKV_PE243_-infected mice was collected from the base of the tail at 4 hours, 48 hours and 144 hours following the appropriate treatments and subsequently centrifuged at 400 g for 30 min for plasma separation. Samples obtained at different periods post infection were titrated using a plaque assay. Vero cells (obtained from ATCC^®^ CCL81™) (African green monkey kidney epithelial cell line) were plated in 24-well plates at 4×10^4^ cells per well in Dulbecco’s modified Eagle’s medium (DMEM) (GIBCO, Thermo Fisher, USA) supplemented with 5% fetal bovine serum (FBS) (GIBCO, Thermo Fisher, USA) and 1% gentamicin (10 μg/ml) (GIBCO, Thermo Fisher, USA) and cultured overnight for complete adhesion at 37°C with 5% CO_2_. Then, the medium was removed, and the cells were washed with 1x PBS and incubated with serial (base 10) dilutions of virus in FBS-free medium. After 90 min of incubation under gentle shaking, the medium was removed, and the cells were washed with 1x PBS and cultured with 1.5% carboxymethylcellulose (CMC) supplemented with 1% FBS (GIBCO, Thermo Fisher, USA). After 5 days, the cells were fixed overnight with 4% formaldehyde and stained with 1% crystal violet in 20% methanol (ISOFAR, Brazil) for 1 hour. Plaques were counted, and the virus yield was calculated and expressed as plaque-forming units per milliliter (PFU/ml).

### 2.6 Histological, immunohistochemistry and TUNEL analyses of the placenta

Placental fragments were fixed overnight and subjected to dehydration (increasing ethanol series; ISOFAR, Brazil), diaphanization with xylol (ISOFAR, Brazil) and paraffin (Histopar, Easypath, Brazil). Sections (5 μm) were prepared using a Rotatory Microtome CUT 5062 (Slee Medical GmbH, Germany) and subjected to immunohistochemistry and TUNEL analyses.

For immunohistochemistry, blocking of endogenous peroxidase was performed with 3% hydrogen peroxide diluted in PBS, followed by microwave antigenic recovery in Tris-EDTA (pH=9) and sodium citrate (pH=6) buffers (15 min for Tris-EDTA buffer and 8 min for citrate buffer). Sections were washed in PBS + 0.2% Tween and exposed to 3% PBS/BSA for 1 hour. Sections were then incubated overnight at 4°C with the following primary antibodies: anti-Ki-67 (1:100 – [M3064]; Spring Bioscience, USA), anti-P-gp (1:500 – Mdr1[sc-55510]; Santa Cruz Biotechnology, USA), anti-Bcrp (1:100 – Bcrp [MAB4146]; Merck Millipore, USA), anti-Abcg1 (1:100 – [PA5-13462]; Thermo Fisher Scientific, USA) or anti-Abca1 (1:100 – [ab18180]; Abcam Plc, UK). The next day, sections were incubated with the biotin-conjugated secondary antibody SPD-060 (Spring Bioscience, USA) for 1 hour at room temperature. Three washes were performed with PBS + 0.2% Tween followed by incubation with streptavidin (SPD-060 - Spring Bioscience, USA) for 30 min. Sections were stained with 3,3-diaminobenzidine (DAB) (SPD-060 - Spring Bioscience, USA), counterstained with hematoxylin (Proquímios, Brazil), dehydrated, diaphanized and mounted with a coverslip and Entellan (Merck, Germany).

For analysis of apoptotic nuclei, terminal deoxynucleotidyl transferase dUTP nick-end labeling (TUNEL) staining was performed using an ApopTag^®^ In Situ Peroxidase Detection Kit (S7100, Merck Millipore, USA) according to the manufacturer’s recommendations and as previously described (36). All negative controls were prepared with omission of the primary antibody.

Image acquisition was performed using a high-resolution Olympus DP72 (Olympus Corporation, Japan) camera coupled to an Olympus BX53 light microscope (Olympus Corporation, Japan). For nuclear quantification of Ki-67 and TUNEL immunolabeling, Stepanizer software (40) was used. For this analysis, we evaluated 15 images from different random fields of the Lz (labyrinth zone) and Jz (junctional zone) for each animal, in a total of five animals from each ICompetent group and three animals from each ICompromised group. A total of 360 digital images (40X) randomly captured per placental region (Lz and Jz) were evaluated in each experimental group. The total number of immunolabeled Ki-67 or TUNEL nuclei in each digital image was normalized by the total image area to obtain an index of the estimated number of proliferative and apoptotic nuclei in the entire histological section analyzed. Analysis was undertaken by two investigators blinded to the treatment.

Quantification of P-gp, Bcrp, Abca1 and Abcg1 staining was performed using the Image-Pro Plus, version 5.0 software (Media Cybernetics, USA) mask tool. The percentage of viable tissue area was considered upon exclusion of negative spaces. A total of 360 digital images (40X) randomly captured per placental region (Lz and Jz) were evaluated in each experimental group. Analysis was undertaken by two investigators blinded to the treatment.

### 2.7 Transmission electron microscopy (TEM)

Sections of the placental Lz and Jz were fixed in paraformaldehyde 4% (Sigma-Aldrich, Brazil) for 48 hours, postfixed with osmium tetroxide (Electron Microscopy Sciences, USA) and potassium ferrocyanide (Electron Microscopy Sciences, USA) for 60 min and dehydrated with an increasing series of acetone (30%, 50%, 70%, 90% and two of 100%) (ISOFAR, Brazil). Sections were subsequently embedded with EPOXI resin (Electron Microscopy Sciences, USA) and acetone (1:2, 1:1 and 2:1, respectively). After polymerization, ultrafine sections (70 nm) were prepared (Leica Microsystems, USA) and collected into 300 mesh copper grids (Electron Microscopy Sciences, USA). Tissue was contrasted with uranyl acetate and lead citrate and visualized using a JEOL JEM-1011 transmission electron microscope (JEOL, Ltd., Akishima, Tokyo, Japan). Digital micrographs were captured using an ORIUS CCD digital camera (Gatan, Inc., Pleasanton, California, EUA) at 6000× magnification. An overall qualitative analysis of the Lz and Jz in different groups was performed by investigating the ultrastructural characteristics of the mitochondria and the ER cisterns. The qualitative evaluation consisted of analyzing disruption of the mitochondrial membranes, mitochondrial morphology, preservation of mitochondrial cristae and matrix intensity (41). Ultrastructural analysis of nuclear morphology and the presence of microvilli in trophoblast sinusoidal giant cells was also undertaken. Analysis of ER cisterns was performed by evaluating the dilation of their lumen (42).

### 2.8 Statistical analysis

GraphPad Prism 8 software (GraphPad Software, Inc., USA) was used for statistical analysis. A D’Agostino & Pearson normality test was used to evaluate normal distribution, and outliers were identified using a Grubbs test. The data are expressed as the mean□±SEM or individual values. One-way ANOVA followed by Tukey’s posttest was used for comparisons between different inbred groups, whereas Student’s t-test or a nonparametric Mann-Whitney test was performed to compare the outbred groups. Differences were considered significant when p<0.05. Pregnancy parameters were evaluated using the mean value of all fetuses and placentae in a litter per dam and not the individual conceptus, i.e., the mean value. In Figures 1 and 2, “n” represents the number of dams. For MET and immunostaining data, placentae closest to the mean weight of all placentae were selected from each litter; “n” represents the number of litters (35–37).

**Figure 1:**
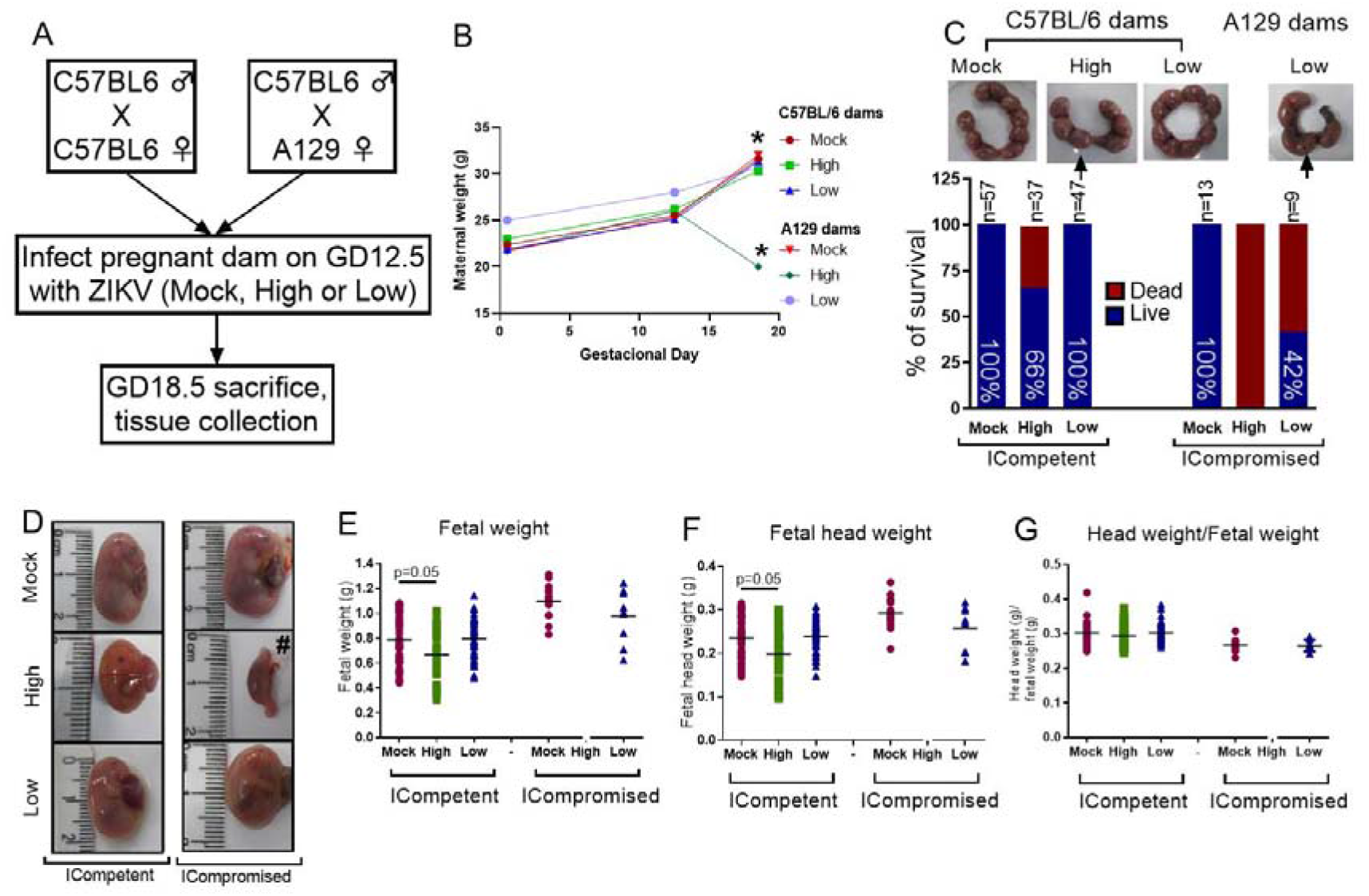
ZIKV infection induced fetal changes during pregnancy. **A)** Experimental design: matings of C57BL/6 dams x C57BL/6 sires (mock n=11 dams; high ZIKV n=15 dams; low ZIKV n=9 dams) and A129 dams x C57BL/6 sires (n=3 dams/group). The mock control consisted of noninfected C6/36 cell supernatants; high ZIKV titers consisted of 5×10^7^ plaque-forming units (PFU) of ZIKV_PE243_ and low ZIKV titers consisted of 10 PFU of ZIKV_PE243_. Fetuses from C57BL/6 dams and sires were termed ICompetent, whereas fetuses from A129 dams and C57BL/6 sires were termed ICompromised. **B)** Maternal weight gain throughout pregnancy, *=p<0.05, one-way ANOVA. **C)** Uterine horn and survival rates following ZIKV exposure (arrows show resorption sites). **D)** Fetal/reabsorption images (#=example of fetal reabsorption). **E)** Fetal weight, **F)** fetal head weight and **G)** fetal head weight/fetal weight ratio. One-way ANOVA followed by Tukey’s posttest was used to assess changes among ICompetent groups, whereas an unpaired Student’s t-test or nonparametric Mann-Whitney test was used to assess differences between ICompromised groups. Values are the mean of individual plotted values.

**Figure 2:**
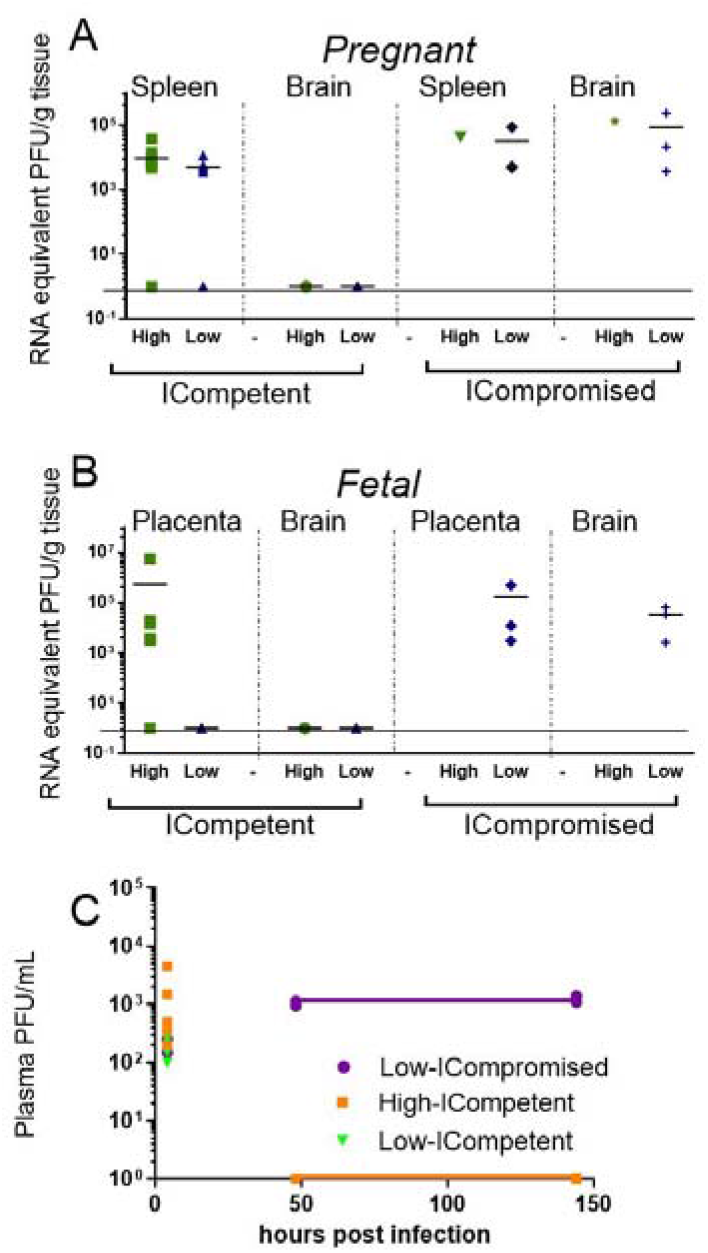
Viral load detection in maternal and fetal tissues after ZIKV infection. Pregnant ICompetent and ICompromised mice were inoculated i.v. with low and high doses of ZIKV. A-B) ZIKV RNA was measured via RT-qPCR in tissue samples obtained from maternal spleen and brain (A) and from placenta and fetal brain (B). C) The presence of ZIKV infectious particles in the plasma of infected dams was evaluated with a plaque assay 4, 48, and 144 hours post infection in ICompetent (mock n=11 dams; high ZIKV n=15 dams; low ZIKV n=9 dams) and ICompromised (mock n=3 dams; high ZIKV n=3 dams; low ZIKV n=3 dams) mice.

## 3. Results

### 3.1 Weight gain during pregnancy is dependent on maternal immune status in ZIKV-infected mice

To determine the effect of ZIKV infection on fetal and placental phenotypes at term (GD18.5), we infected ICompetent C57BL/6 and immunocompromised (ICompromised) A129 mice with ZIKV at GD12.5 (Figure 1A). Given the very distinct susceptibility of C57BL/6 and A129 mice to ZIKV, systemic infection models were established by injecting high (5×10 PFU) and low (10 PFU) virus inoculum titers. As shown in Figure 1B, ICompetent C57BL/6 mice in all groups and ICompromised A129 dams inoculated with mock and low ZIKV titers exhibited higher maternal weight at GD18.5 than at GD12.5 and GD0.5 (p<0.05). On the other hand, ICompromised A129 mice presented significant weight loss at GD18.5, despite showing an increase at GD12.5 in relation to GD0.5 (Figure 1B).

### 3.2 Immunocompetent and immunodeficient mice have distinct term placental and fetal phenotypes in response to high and low ZIKV titer challenges in mid-pregnancy

The fetuses from C57BL/6 dams and sires were called ICompetent. The fetuses from the mating of A129 dams and C57BL/6 sires were called ICompromised. High-ZIKV ICompetent mice exhibited 34% fetal loss, whereas high-ZIKV-A129 mice had 100% fetal loss (Figure 1C). In the low-ZIKV groups, C57BL/6 mice had no (0%) fetal death, while A129 mice exhibited a 42% fetal death rate (Figure 1C). Fetal and fetal head sizes were decreased in A129 mice compared to those in C57BL/6 dams infected with the high ZIKV titer (p=0.05; Figure 1 D-G). However, no changes in fetal weight or fetal head size were observed when the mice were infected with the low ZIKV titer (Figure 1 D-G).

### 3.3 ZIKV is detected in the fetal brain of ICompromised, but not ICompetent mice

ZIKV RNA was detected in the spleens of pregnant ICompetent C57BL/6 mice inoculated with the highest ZIKV titer, confirming acute systemic infection. Viral RNA was also detected in the majority of the placentae of those mice (Figure 2A-B) but not in the maternal and fetal C57BL/6 brains (Figure 2A-B), suggesting that the virus was not transmitted to the fetuses. Although low ZIKV inoculation resulted in virus detection in the spleens of pregnant C57BL/6 mice, infection was not evidenced in the placentae or fetal brains. In contrast, ZIKV RNA was detected in all analyzed organs from ICompromised A129 dams, including the maternal brain and spleen and the placenta and fetal brain (Figure 2A-B). Viremia in maternal plasma was evaluated at 4, 48 and 144 hours after infection. Within 4 hours, the presence of the virus was verified in the serum (high ICompetent=637.5 PFU/mL, low ICompetent=740 PFU/mL and low ICompromised=325 PFU/mL), indicating that the virus was correctly inoculated. Afterwards, ZIKV RNA was detected in ICompromised dams at 48 and 144 hours post inoculation but not in ICompetent dams (Figure 2C).

### 3.4 ZIKV infection induces distinct systemic and placental inflammatory responses in ICompetent and ICompromised mice

The maternal serum and placental protein levels of specific cytokines and chemokines related to fetal death and preterm delivery (43–45) were evaluated to probe whether midgestation ZIKV infection would induce a maternal inflammatory response at term in our two distinct models. Since A129 infected with 10^7^ ZIKV-PFU showed 100% fetal loss, we proceeded using 10^7^ PFU inoculation in C57BL/6 mice and 10^3^ PFU inoculation in both C57BL/6 and A129 mice.

CCL2 was elevated in the serum of low-ZIKV ICompetent mice, but no other alteration was systemically detected in any ICompetent mice at this time point (Figure 3A-B). On the other hand, ICompromised dams showed significantly increased CXCL1 and IL-6 levels in the serum and a strong trend for an enhancement in CCL2 (Figure 3C). Analysis of cytokine and chemokine expression in the placenta demonstrated that the CXCL1 and CCL2 chemokines were also upregulated in ICompromised but not ICompetent mice (Figure 3D-F). Surprisingly, IL-6 protein expression was augmented in some of the low-ZIKV ICompetent mice (58%; p<0.05) but not in high-ZIKV mice or ICompromised mice (Figure 3D-F).

**Figure 3:**
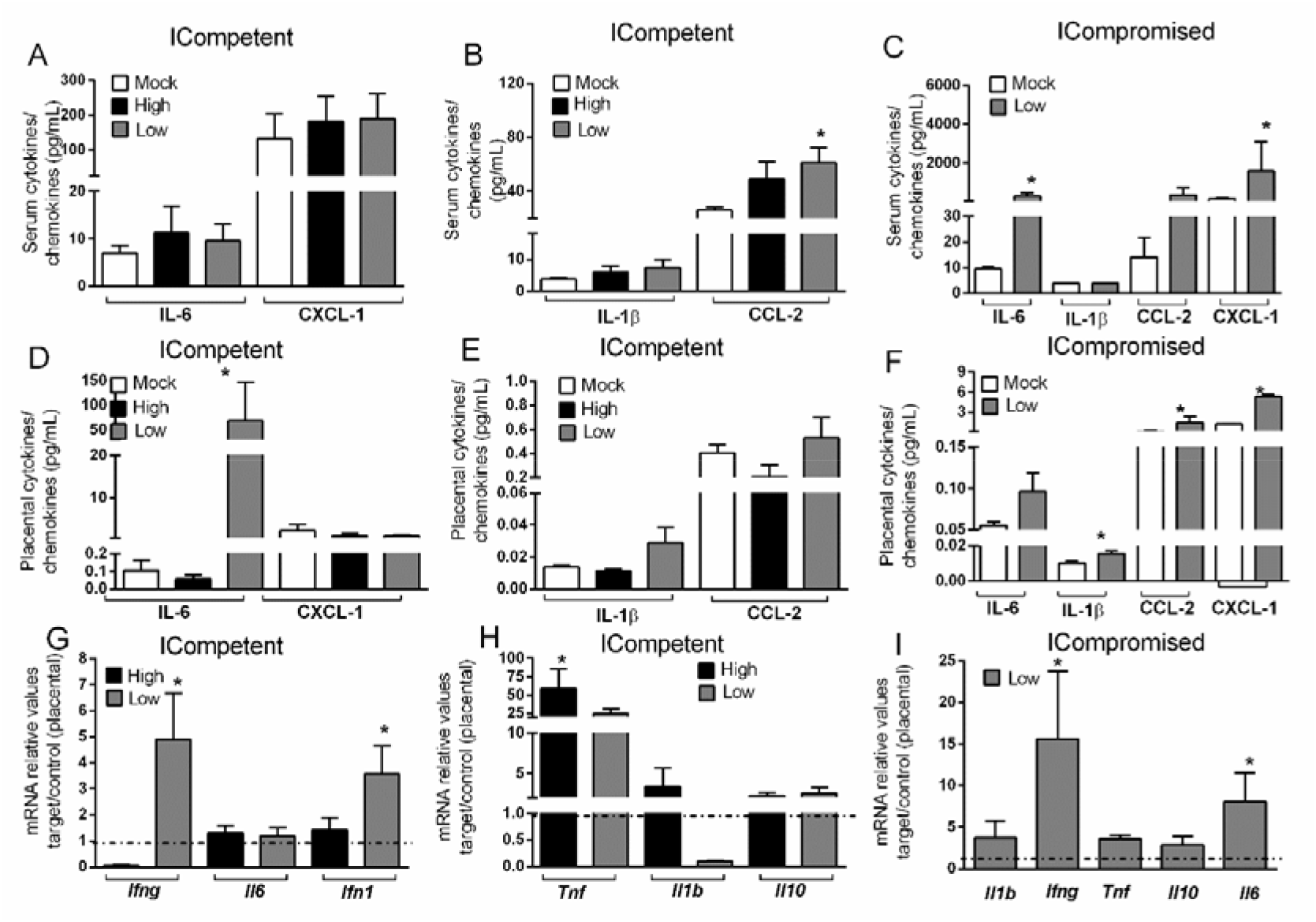
ZIKV during pregnancy promotes an inflammatory response in the maternal serum and in the placenta. **A)** Levels of IL-1β, IL-6, CCL-2 and CXCL-1 in the maternal serum (A-C) and placenta (D-F) at GD18.5 in matings of C57BL/6 dams x C57BL/6 sires (ICompetent fetuses - mock n=11 dams; high ZIKV n=15 dams; low ZIKV n=9 dams) and A129 dams x C57BL/6 sires (ICompromised fetuses n=3 dams/group). Placental mRNA expression (G-I) of *Ifng, Ifn1, Il1b, Il6, Il10 and Tnf*. Broken lines show the expression levels in both lineages in the mock group. One-way ANOVA followed by Tukey’s posttest was used to assess changes among ICompetent groups, whereas an unpaired Student’s t-test or nonparametric Mann-Whitney test was used to assess differences between ICompromised groups. The values are expressed as the mean ± SEM.

We also assessed the placental mRNA expression of a range of cytokines related to placental infective responses: *Ifng, Il6, Ifn1, Tnf, Il1b and Il10*. All the infected mice showed a significant increase in *Tnf* mRNA expression, with higher levels detected in ICompetent mice (Figure 3G-I). The low-ZIKV ICompetent mice presented modest but significant *Ifn1* expression, which was not detected in the high-ZIKV group (Figure 3G). Additionally, both low-ZIKV-infected groups (ICompetent and ICompromised) presented increased *Ifng* mRNA expression (Figure 3G and 3I, p< 0.05). Interestingly, placental *Il6* mRNA levels were only elevated in ICompromised pregnancies compared to mock pregnancies (p=0.05) (Figure 3I). *Il1b* and *Il10* remained unchanged in all groups analyzed (Figure 3 G-I).

### 3.5 ZIKV affects placental proliferation, apoptosis and ultrastructure in a viral load- and maternal immune status-dependent manner

We did not observe changes in placental weight or in the fetal:placental weight ratio in any of the groups investigated (data not shown). However, since we detected the presence of ZIKV RNA in the placentas, we investigated the cellular proliferation (Ki-67^+^ cells) and apoptotic ratio in the Lz and Jz of the mouse placenta. Increased Ki-67 staining was observed in the Lz of high- and low-ZIKV-treated ICompetent animals compared to mock animals (Figure 4A-D, p=0.005 and p=0.015), whereas the apoptotic ratio in the Lz was increased in high-ZIKV dams and decreased in low-ZIKV dams (p=0.01, Figure 4E-H). Although no differences were observed in Ki67 staining (Figure 5A-D), a similar apoptotic pattern was detected in the Jz of ICompetent pregnancies (p=0.008 and p=0.004, respectively; Figure 5E-H). In contrast, in ICompromised dams, Lz Ki-67 staining was decreased (p=0.001; Figure 4I-L), while the apoptotic reaction was increased in ZIKV-infected animals (p=0.05; Figure 4M-P). Jz from ICompromised offspring exhibited increased Ki-67 labeling and no differences in the apoptotic reaction (p=0.001; Figure 5 I-L and Figure 5M-P).

**Figure 4:**
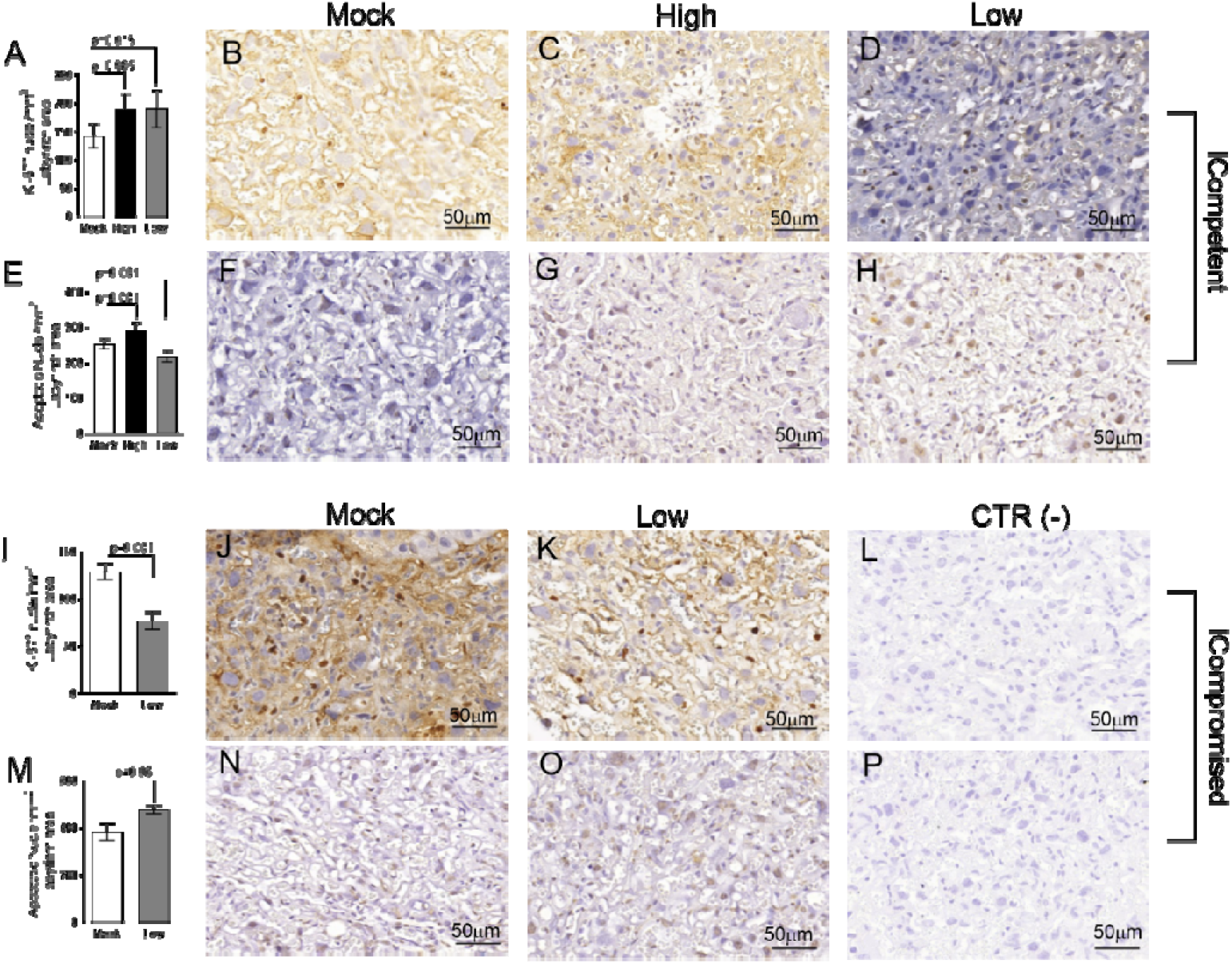
Labyrinthine remodeling in ICompetent and ICompromised placentae is affected by gestational ZIKV infection. A total of 180 digital images (40X) randomly captured from the whole labyrinth zone (Lz) of each placenta per dam were evaluated. Immunolabeled nuclei from each digital image were quantified and normalized by the total digital image area to obtain an index of the estimated number of proliferative and apoptotic nuclei in the entire histological section. (**A and I**) Quantification and (**B-D and J-K**) representative photomicrographs of Ki-67^+^ stained nuclei in the Lz of ICompetent (n=6 placentae from 6 independent dams/group) and ICompromised (n=3 placentae from 3 independent dams/group) placentae, respectively. (**E and M**) Quantification and (**F-H and N-O**) representative photomicrographs of apoptotic nuclei (TUNEL) in the Lz of ICompetent (n=5 placentae from 5 independent dams/group) and ICompromised (n=3 placentae from 3 independent dams/group) placentae, respectively. (**L and P**) Negative controls. One-way ANOVA followed by Tukey’s posttest was used to assess changes among ICompetent groups, whereas an unpaired Student’s t-test or nonparametric Mann-Whitney test was used to assess differences between ICompromised groups. The values are expressed as the mean ± SEM. Images were captured at 40X. Scale bar=50 μm.

**Figure 5:**
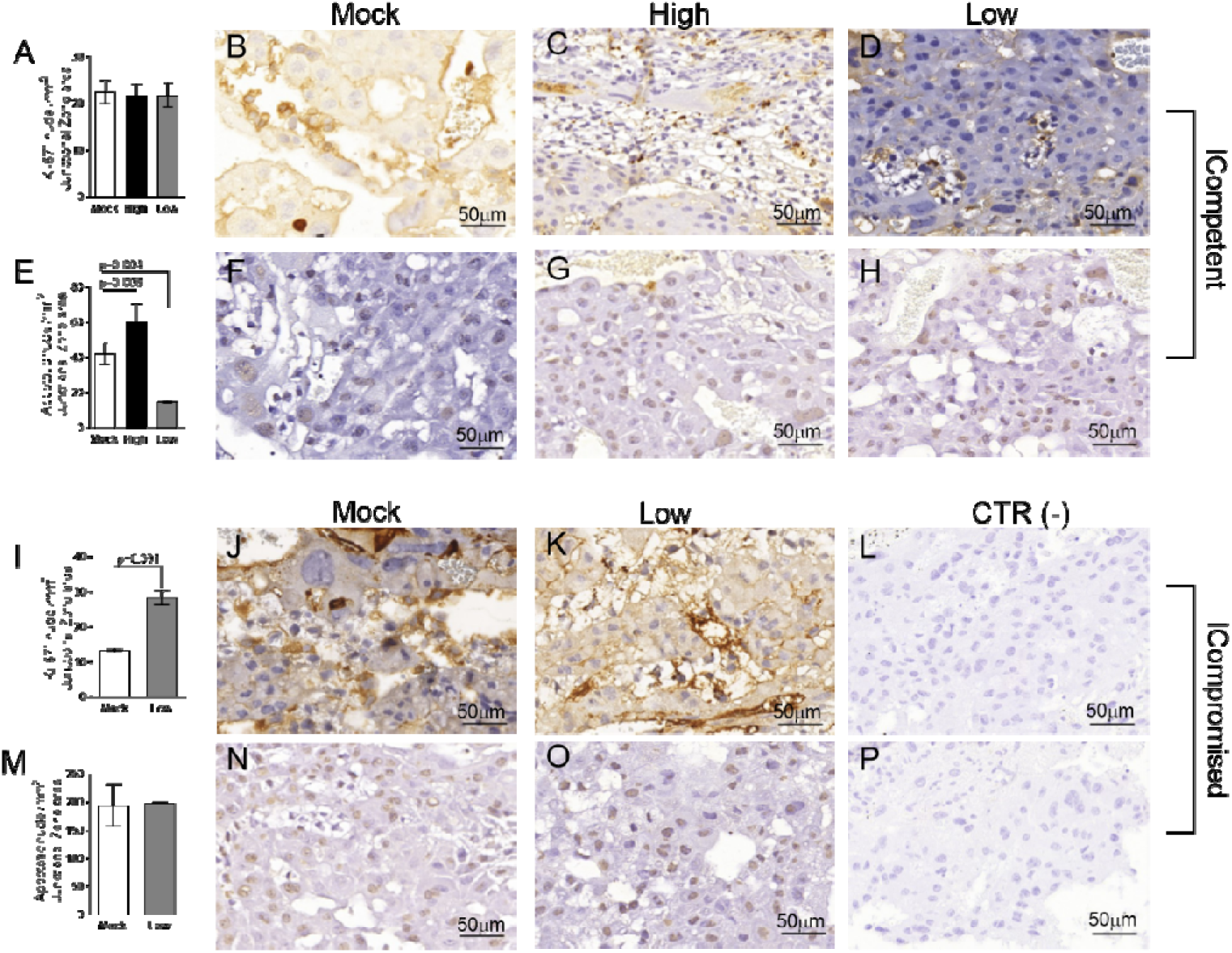
Junctional zone remodeling in ICompetent and ICompromised placentae is affected by gestational ZIKV infection. A total of 180 digital images (40X) randomly captured from the whole junctional zone (Jz) of each placenta per dam were evaluated. Immunolabeled nuclei from each digital image were quantified and normalized by the total digital image area to obtain an index of the estimated number of proliferative and apoptotic nuclei in the entire histological section. (**A and I**) Quantification and (**B-D and J-K**) representative photomicrographs of Ki-67^+^ stained nuclei in the Jz of ICompetent (n=6 placentae from 6 independent dams/group) and ICompromised (n=3 placentae from 3 independent dams/group) placentae, respectively. (**E and M**) Quantification and (**F-H and N-O**) representative photomicrographs of apoptotic nuclei (TUNEL) in the Jz of ICompetent (n=5 placentae from 5 independent dams/group) and ICompromised (n=3 placentae from 3 independent dams/group) placentae, respectively. (**L and P**) Negative controls. One-way ANOVA followed by Tukey’s posttest was used to assess changes among ICompetent groups, whereas an unpaired Student’s t-test or nonparametric Mann-Whitney test was used to assess differences between ICompromised groups. The values are expressed as the mean ± SEM. Images were captured at 40X. Scale bar=50 μm.

### 3.6 Placental ultrastructure is differently impacted by high- and low-titer ZIKV infection in ICompetent and ICompromised strains

Lz ultrastructural analyses of ICompetent and ICompromised-mock animals detected sinusoidal trophoblastic giant cells exhibiting regular microvilli, euchromatic nuclei, preserved mitochondrial ultrastructure and regular narrow ER cisternae (Figure 6A and 6B). In sharp contrast, high-ZIKV ICompetent (Figure 6C) infected placentae showed fewer villi in the sinusoidal giant trophoblastic cells, degenerated mitochondria, granular ER with dilated cisterns and euchromatic nuclei. The sinusoidal giant trophoblastic cells in the low-ZIKV ICompetent mice (Figure 6D) also had fewer villi and degenerated mitochondria than those in the mock placentae, but no effect on the ER or euchromatic nuclei observed. Low-ZIKV ICompromised infected placentae (Figure 6E) showed fewer villi in the sinusoidal giant trophoblastic cells, degenerated mitochondria, granular ER with dilated cisterns and euchromatic nuclei.

**Figure 6:**
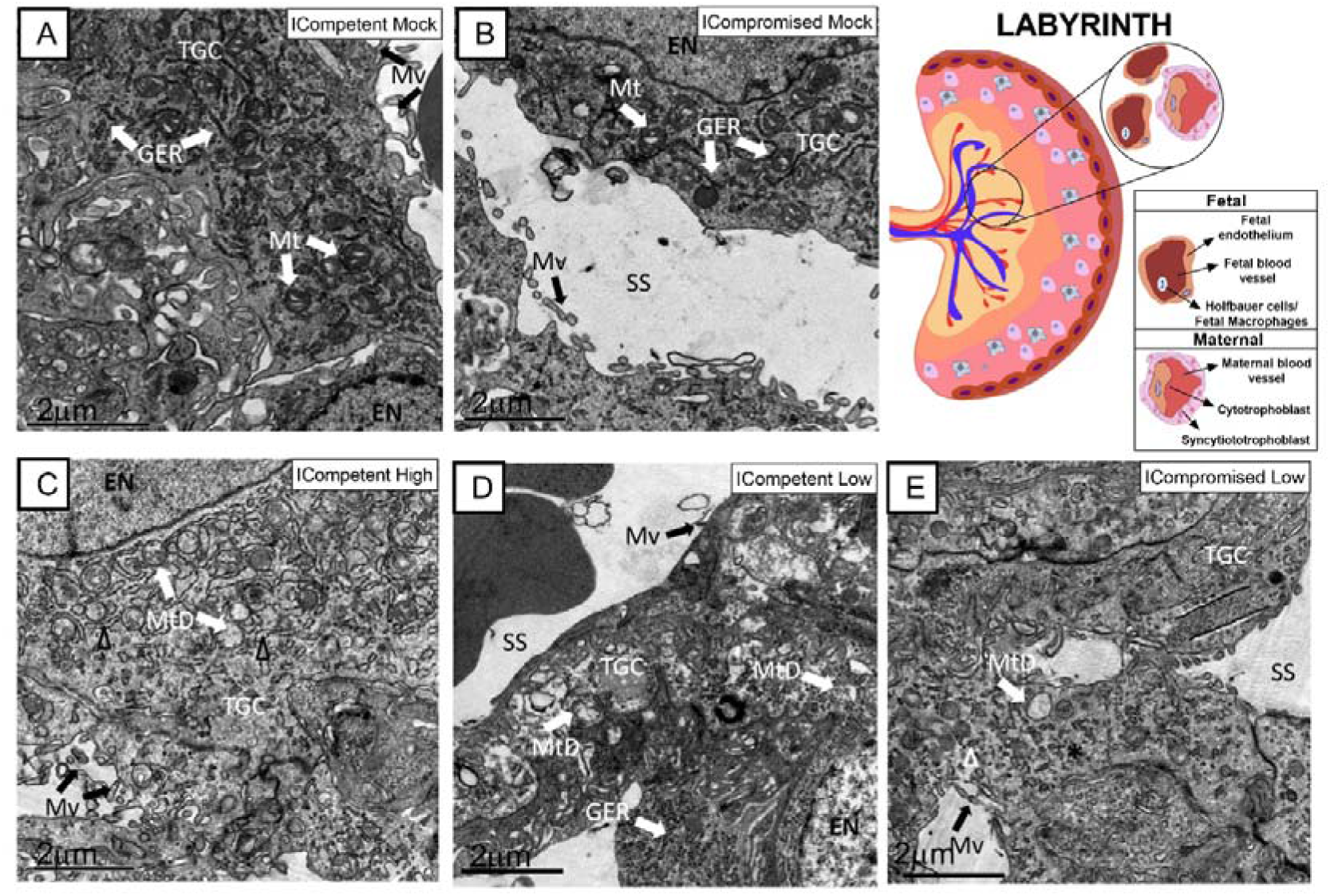
Associated ultrastructural changes in the placental Lz after ZIKV infection. Transmission electron photomicrographs of ICompetent mock (A), ICompromised mock (B), ICompetent high (C), ICompetent low (D) and ICompromised low (E) groups (n=5/group). We observed dilatation in the ER cisterns of ICompetent high placentas. Additionally, there was a reduction in the microvilli in both the ICompetent high and ICompetent low placentas. In the ICompromised low group, we found fragmented ER and microvillus reduction. All infected groups showed degenerate mitochondria. GER=granular endoplasmic reticulum; Δ=dilated granular endoplasmic reticulum; *=fragmented granular endoplasmic reticulum; Mt=mitochondria; MtD=degenerate mitochondria; Mv=microvilli; EN=euchromatic nuclei; SS=sinusoidal space; TGC=trophoblastic giant cell. Scale bar=2 μm.

The Jz of mock ICompetent and ICompromised placentae (Figure 7A and 7B) exhibited euchromatic nuclei, with evident heterochromatin, preserved mitochondria and narrow cisternae in a granular ER. High-ZIKV ICompetent Jz had degenerated mitochondria, granular ER with dilated cisterns and euchromatic nuclei (Figure 7C). Low-ZIKV ICompetent (Figure 7D) placentae exhibited euchromatic nuclei with evident heterochromatin, preserved mitochondria and narrow cisternae in a granular ER, whereas low-ZIKV ICompromised placentae had degenerated mitochondria, granular ER with dilated cisterns and euchromatic nuclei (Figure 7E).

**Figure 7:**
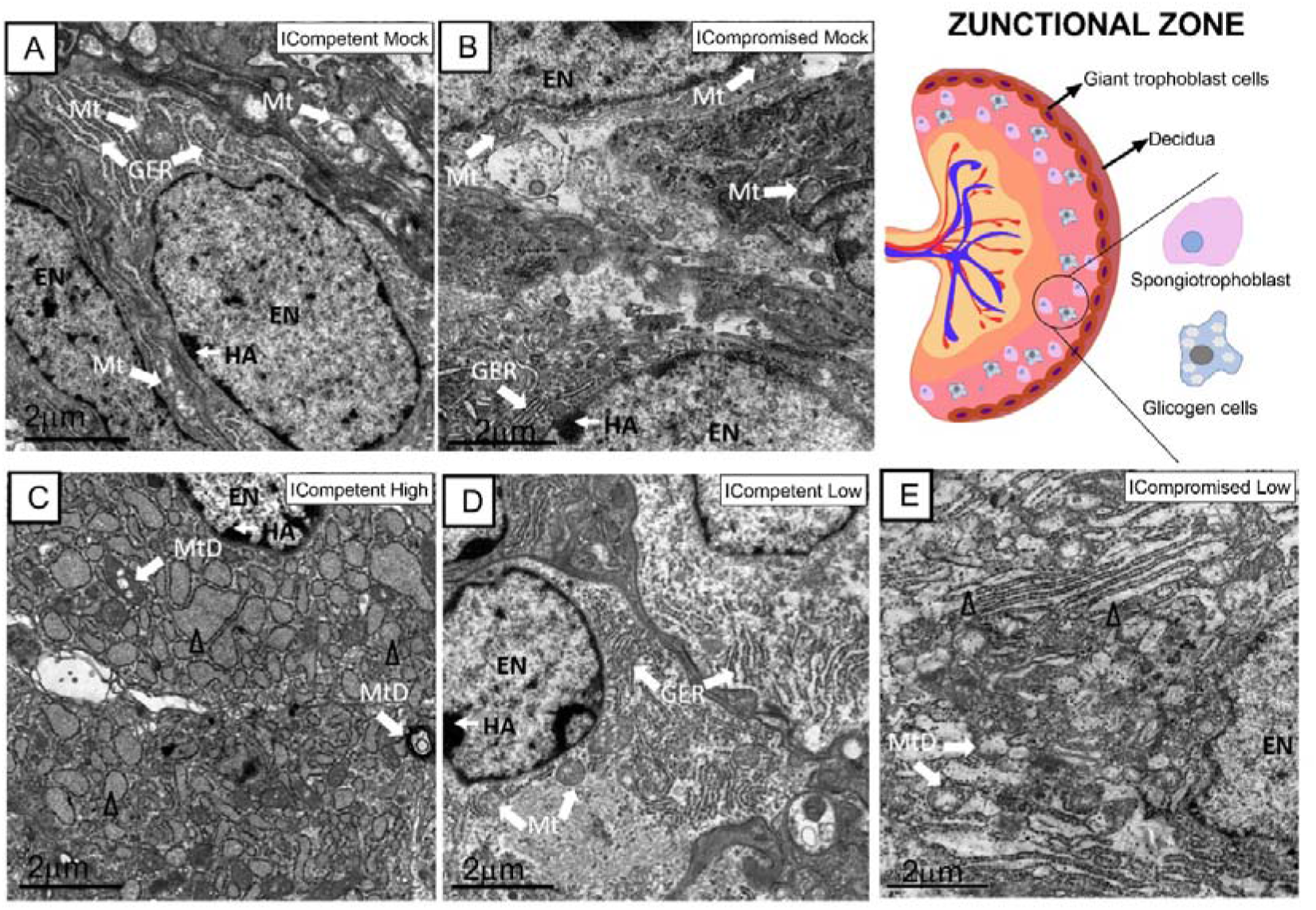
Associated ultrastructural changes in the placental Jz after ZIKV infection. Transmission electron photomicrographs of ICompetent mock (A), Compromised mock (B), ICompetent high (C), ICompetent low (D) and ICompromised low (E) groups (n=5/group). We found deteriorating mitochondria and dilated reticulum endoplasmic cisterns in both the high ICompetent and low ICompromised groups. GER=granular endoplasmic reticulum; Δ=dilated granular endoplasmic reticulum; Mt=mitochondria; MtD=degenerate mitochondria; Mv=microvilli; EN=euchromatic nuclei; SS=sinusoidal space; TGC=trophoblastic giant cell; HA=heterochromatin area. Scale bar=2 μm.

### 3.7 ZIKV differentially affects placental expression of drug and lipid ABC transporter systems

Evaluation of key ABC transporters in the Lz of mock and ZIKV-infected ICompetent and ICompromised placentae revealed that immunolabeling of the drug P-gp and Bcrp efflux transporter systems was primarily present at the cellular membranes of the sinusoidal trophoblastic giant cells, with diffuse cytoplasmic Bcrp staining. Labeling of the Abca1 and Abcg1 lipid efflux transporters was moderately and heterogeneously distributed within the Lz. Less Lz-P-gp was observed in ICompetent mice infected with both high- and low-ZIKV infective regimens than in mock-treated animals (p=0.001 and p=0.002, respectively; Figure 8A-E), whereas reduced Bcrp and Abca1 staining was observed in high-ZIKV-infected mice (p=0.003 and p=0.004, Figure 8F-J and Figure 8K-O, respectively). No changes in Abcg1 were observed in any of the ICompetent experimental groups (Figure 8P-T). P-gp, Bcrp and Abca1 transporter immunostaining was downregulated in ICompromised low ZIKV-treated animals (p=0.001, p=0.05 and p=0.05, Figure 9A-D, Figure 9E-H and Figure 9I-L, respectively). No changes in Abcg1 were observed in any of the ICompromised experimental groups (Figure 9M-P).

**Figure 8:**
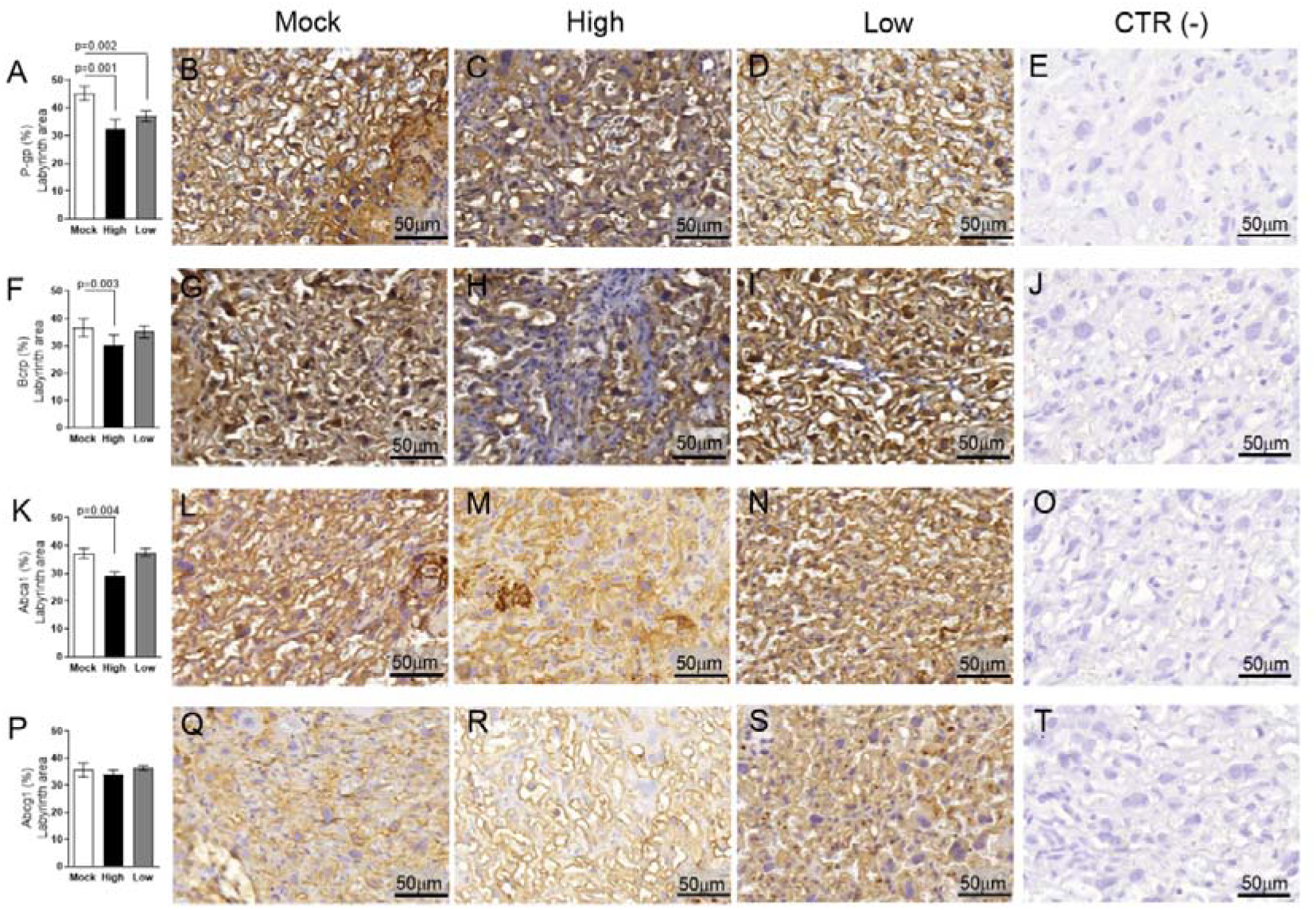
ZIKV infection decreases P-gp, Bcrp and Abca1 expression in the placental Lz of infected mice in the ICompetent groups. A total of 180 digital images (40X) randomly captured from the whole labyrinth zone (Lz) of each placenta per dam were evaluated. Immunolabeling in each digital image was quantified by calculating the percentage area of the total stained labyrinthine tissue after exclusion of the total negative space. (**A**) Quantification and (**B-D**) representative photomicrographs of P-gp staining in the Lz of ICompetent (n=6 placentae from 6 independent dams/group) placenta. (**F**) Quantification and (**G-I**) representative photomicrographs of Bcrp staining in the Lz of ICompetent (n=6 placentae from 6 independent dams/group) placenta. (**K**) Quantification and (**L-N**) representative photomicrographs of Abca1 staining in the Lz of ICompetent (n=6 placentae from 6 independent dams/group) placenta. (**P**) Quantification and (**Q-S**) representative photomicrographs of Abcg1 staining in the Lz of ICompetent (n=6 placentae from 6 independent dams/group) placenta. (**E, J, O, T**) Negative controls. One-way ANOVA followed by Tukey’s post-test. The values are expressed as the mean ± SEM. Images were captured at 40X. Scale bar=50 μm.

**Figure 9:**
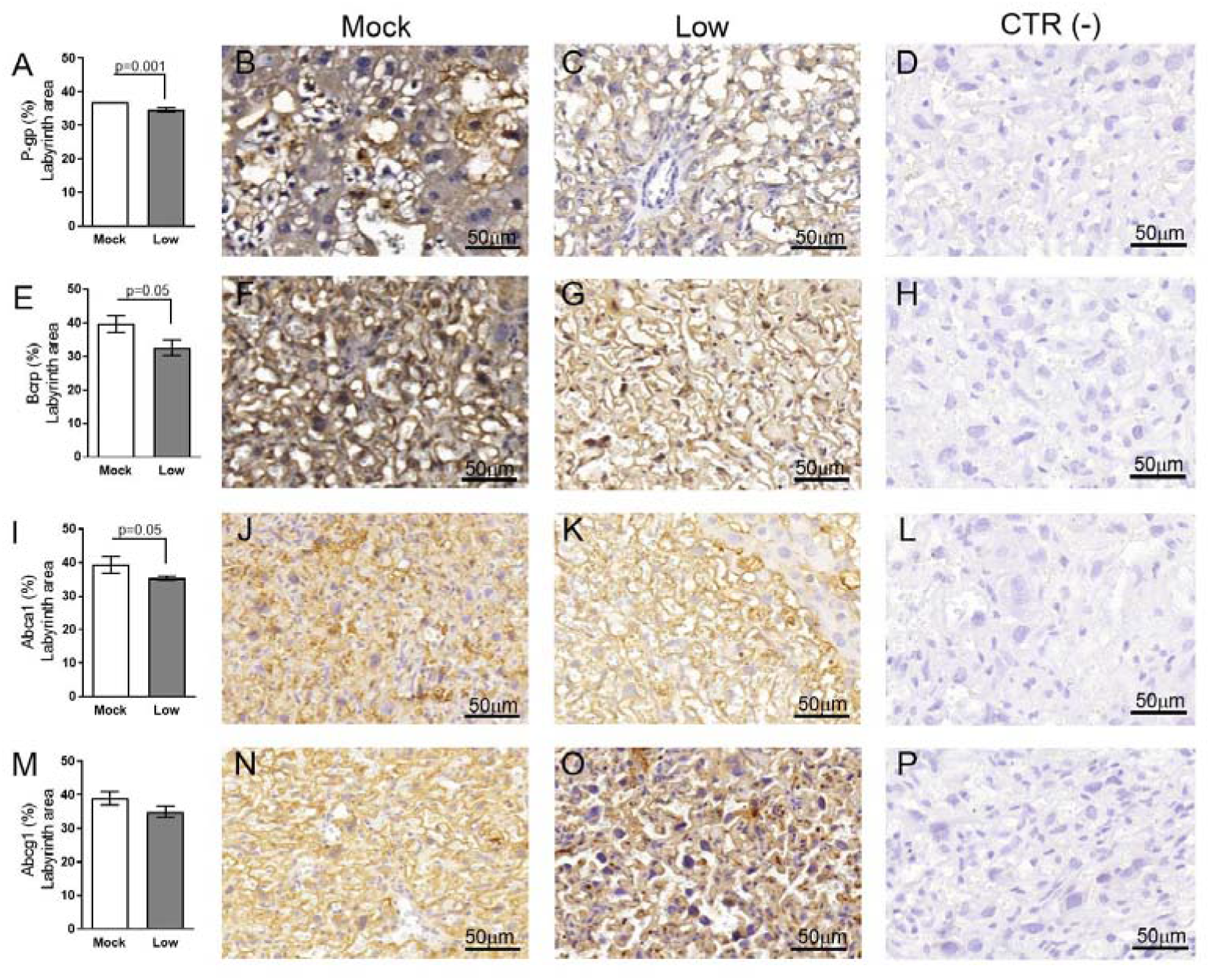
ZIKV infection decreases P-gp, Bcrp and Abca1 protein expression in the placental Lz of infected mice in the ICompromised groups. A total of 180 digital images (40X) randomly captured from the whole labyrinth zone (Lz) of each placenta per dam were evaluated. Immunolabeling in each digital image was quantified by calculating the percentage area of the total stained labyrinth zone tissue after exclusion of the total negative space. (**A**) Quantification and (**B-C**) representative photomicrographs of P-gp staining in the Lz of ICompromised (n=3 placentae from 3 independent dams/group) placenta. (**E**) Quantification and (**F-G**) representative photomicrographs of Bcrp staining in the Lz of ICompromised (n=3 placentae from 3 independent dams/group) placenta. (**I**) Quantification and (**J-K**) representative photomicrographs of Abca1 staining in the Lz of ICompromised (n=3 placentae from 3 independent dams/group) placenta (**M**) quantification and (**N-O**) representative photomicrographs of Abcg1 staining in the Lz of ICompromised (n=3 placentae from 3 independent dams/group) placenta. (**D, H, L, P**) Negative controls. Unpaired Student’s t-test or nonparametric Mann-Whitney test was used to assess differences between ICompromised groups. The values are expressed as the mean ± SEM. Images were captured at 40X. Scale bar=50 μm.

Next, the impact of ZIKV on ABC transporters in the Jz layer (structural and endocrine layers of the mouse placenta) was assessed. P-gp and Bcrp were predominantly localized at the cellular membranes of spongiotrophoblast cells, whereas Abca1 and Abcg1 exhibited membrane and cytoplasmic staining. P-gp staining was decreased in Jz cells from the low-ZIKV ICompromised placentae (p=0.006), with no other alterations observed (Figure 10A-T and Figure 11A-P).

**Figure 10:**
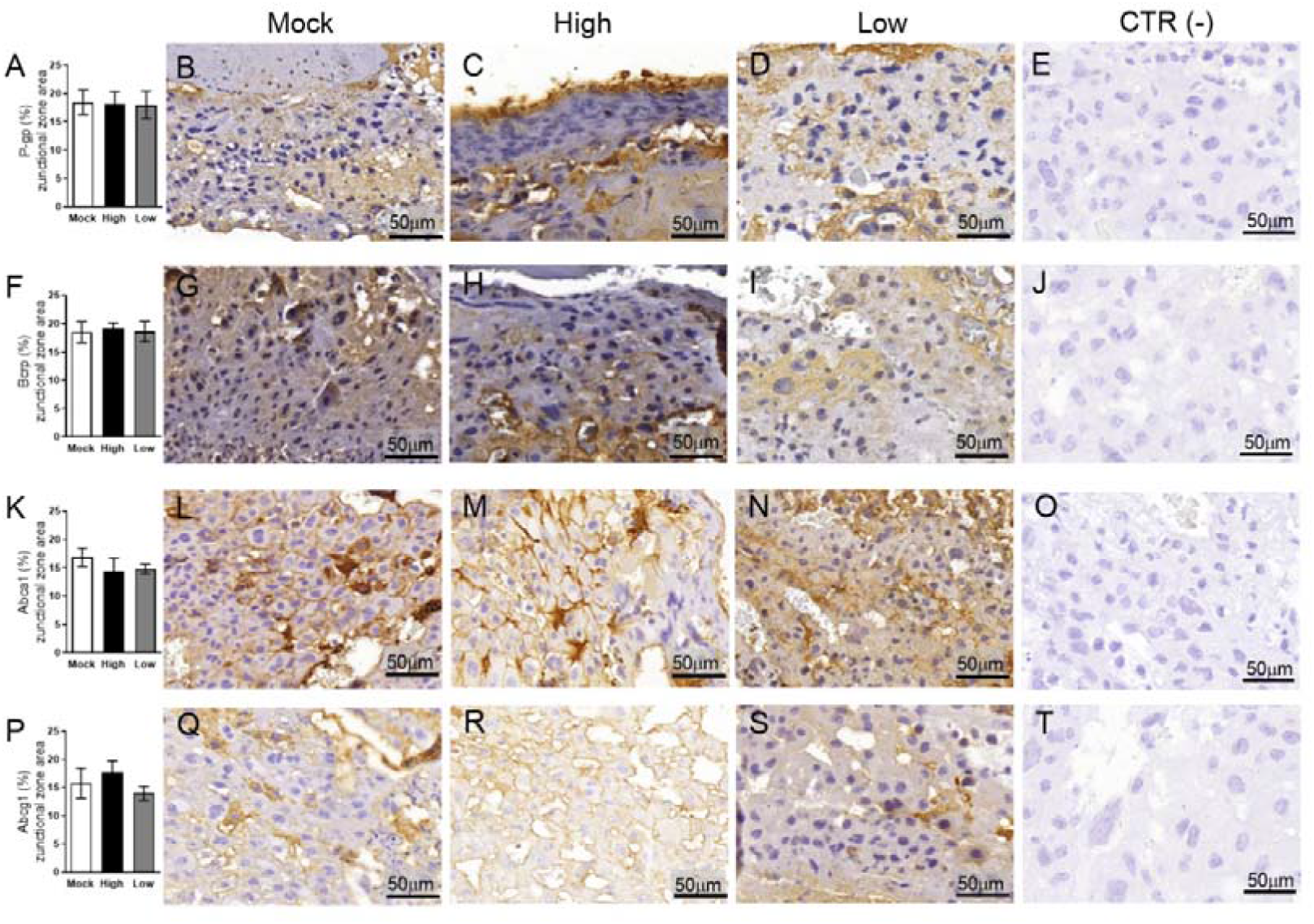
ZIKV infection did not impact P-gp, Bcrp, Abca1 or Abcg1 protein expression in the placental Jz of infected mice in the ICompetent groups. A total of 180 digital images (40X) randomly captured from the whole junctional zone (Jz) of each placenta per dam were evaluated. Immunolabeling in each digital image was quantified by calculating the percentage area of the total stained junctional zone tissue after exclusion of the total negative space. (**A**) Quantification and (**B-D**) representative photomicrographs of P-gp staining in the Jz of ICompetent (n=6 placentae from 6 independent dams/group) placenta. (**F**) Quantification and (**G-I**) representative photomicrographs of Bcrp staining in the Jz of ICompetent (n=6 placentae from 6 independent dams/group) placenta. (**K**) Quantification and (**L-N**) representative photomicrographs of Abca1 staining in the Jz of ICompetent (n=6 placentae from 6 independent dams/group) placenta. (**P**) Quantification and (**Q-S**) representative photomicrographs of Abcg1 staining in the Jz of ICompetent (n=6 placentae from 6 independent dams/group) placenta. (**E, J, O, T**) Negative controls. One-way ANOVA followed by Tukey’s post-test. The values are expressed as the mean ± SEM. Images were captured at 40X. Scale bar=50 μm.

**Figure 11:**
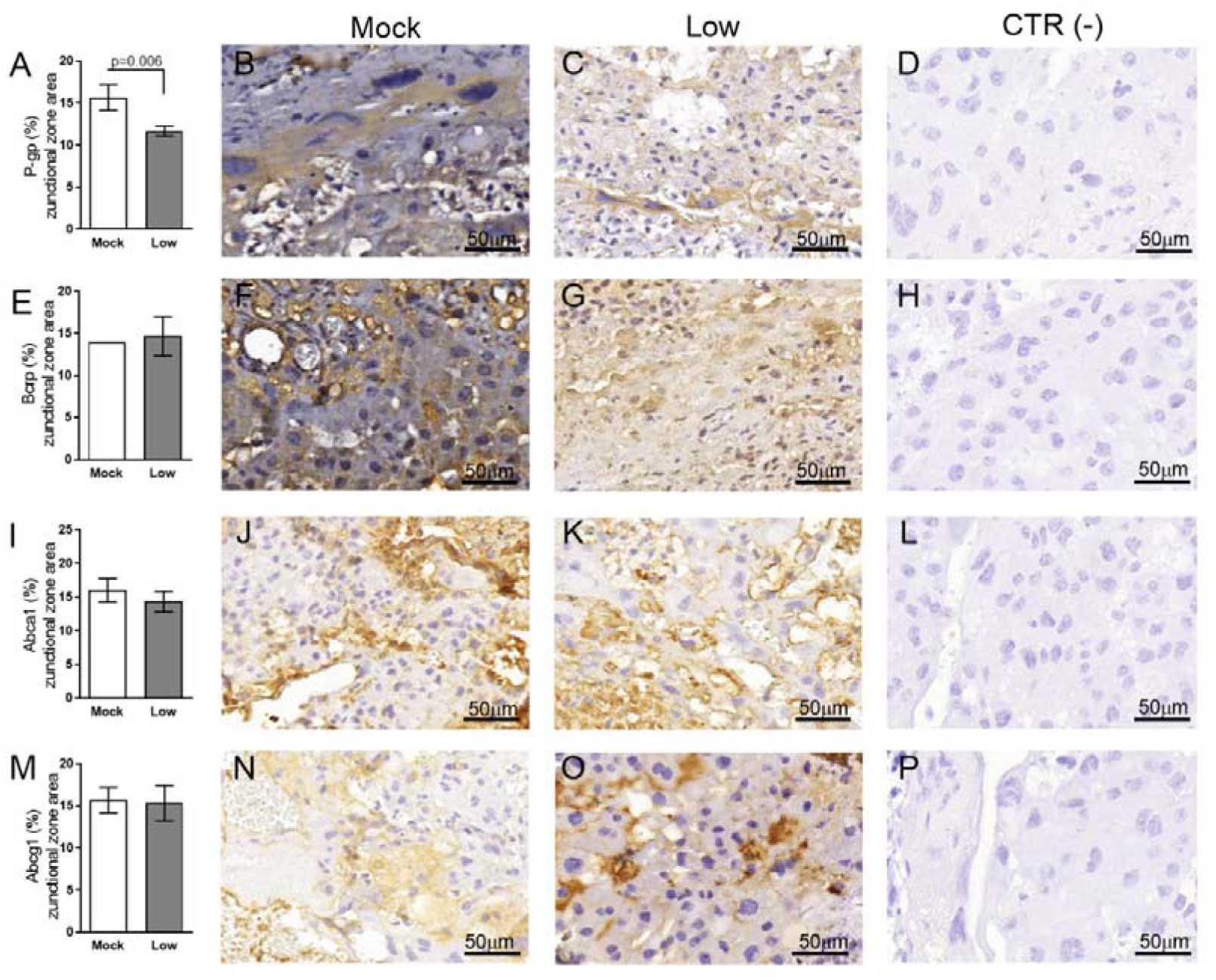
ZIKV infection decreases Bcrp protein expression in the placental Jz of infected mice in the ICompromised group. A total of 180 digital images (40X) randomly captured from the whole junctional zone (Jz) of each placenta per dam were evaluated. Immunolabeling in each digital image was quantified by calculating the percentage area of the total stained junctional zone tissue after exclusion of the total negative space. (**A**) Quantification and (**B-C**) representative photomicrographs of P-gp staining in the Jz of ICompromised (n=3 placentae from 3 independent dams/group) placenta. (**E**) Quantification and (**F-G**) representative photomicrographs of Bcrp staining in the Jz of ICompromised (n=3 placentae from 3 independent dams/group) placenta. (**I**) Quantification and (**J-K**) representative photomicrographs of Abca1 staining in the Jz of ICompromised (n=3 placentae from 3 independent dams/group) placenta. (**M**) Quantification and (**N-O**) representative photomicrographs of Abcg1 staining in the Jz of ICompromised (n=3 placentae from 3 independent dams/group) placenta. (**D, H, L, P**) Negative controls. Unpaired Student’s t-test or nonparametric Mann-Whitney test was used to assess differences between ICompromised groups. The values are expressed as the mean ± SEM. Images were captured at 40X. Scale bar=50 μm.

## 4. Discussion

In this study, we investigated several fetal and placental features at term (GD18.5) in ICompetent (C57BL/6) and ICompromised (A129) mice exposed to ZIKV at mid-pregnancy (GD12.5). Fetal survival rates, systemic and placental inflammatory responses, placental ultrastructure and cell turnover, as well as the expression of key drug (P-gp and Bcrp) and lipid (Abca1) efflux transporter systems in the placenta, were consistently impacted by ZIKV in both strains. The magnitude of the effects was clearly related to the infective titer (high and low) of ZIKV and maternal immune status (ICompetent-C57BL/6 x and ICompromised-A129), and fetal alterations were not exclusively dependent on virus detection in the fetuses.

Infection of ICompetent mice with ZIKV did not result in viremia in the initial postinoculation phase, although viral RNA was detected in the maternal spleen in both the high- and low-ZIKV groups, confirming systemic infection. This is consistent with a previous report (46). Our data demonstrate that pregnant ICompetent C57BL/6 mice were more susceptible to high ZIKV titers than to low ZIKV infective. Since viral RNA was only detected in the placentae of high ZIKV-infected mice, fetal survival rates and weights were impacted to a greater extent in those mice. Strikingly, even though the virus was not present in the fetal brain (at least at term), fetal and fetal head weights were lower in mice subjected to the high-ZIKV titer regimen, suggesting that high infective viral load in mid-pregnancy, even in ICompetent individuals, can induce IUGR and lower fetal head weight despite a lack of transmission to the fetal brain (47). On the other hand, ICompromised placentae and fetal brains had detectable viral transcripts, with no changes in weight, which is consistent with previous data (19). In fact, in our models, the presence (ICompromised) or absence (ICompetent) of the virus in the fetal brain did not correspond to fetal head size (decreased in only high ICompetent). The data from ICompetent and ICompromised placentae demonstrate how important the maternal immunological status is to control viremia, fetal survival and accessibility of the virus to the fetal brain. The reason for the reduction in fetal brain size in C57BL/6 mice in the absence of fetal brain infection requires further investigation. It is possible that fetal brains in the ICompetent mice may have been exposed to ZIKV earlier in pregnancy, when viremia was present in the maternal blood, and this may have severely compromised brain development. Of note, one limitation of our study is that we measured fetal head weight instead of cortical thickness. Future studies should investigate whether high- and low-ZIKV exposure alters cortical thickness in ICompetent and ICompromised offspring.

A distinct inflammatory profile was also detected in the three analyzed groups. At the protein level, low-ZIKV-ICompromised dams exhibited increased maternal IL-6 and CXCL-1 and placental CCL-2 and CXCL-1, whereas low-ZIKV-ICompetent dams had increased maternal CCL2 and placental IL-6 levels. CCCL-2 and CXCL-1 are related to fetal death and preterm delivery (43–45) and could be associated with pronounced fetal injury detected upon ICompromised pregnancy. In addition, IL-6 was previously demonstrated (48) to be related to fetal response syndrome, characterized by activation of the fetal immune system. This syndrome is known to increase fetal morbidity and affect several organs, such as the adrenal gland, brain and heart (32,48–51). At the mRNA level, *Il6* expression was only detected in ICompromised placentae at term and may indicate a sustained harmful response in these mice until term. The IFN signaling pathway may be triggered by ZIKV (17) and is one of the key mechanisms of host defense and a viral target for immune evasion (20), but we only detected a slight increase in *Ifn1* in low-ZIKV ICompetent mice at term. However, we cannot rule out the possibility that these cytokines might have been produced earlier. Our findings showed that *Ifng* expression was significantly enhanced in both ICompetent and ICompromised low-ZIKV-derived placentae but not in high-ZIKV-infected mice. Although we could not assess cytokine expression in high-ZIKV ICompromised placentas, one may extrapolate that low-ZIKV infection could result in stimulation of *Ifng* producing cells, which has been previously shown to be protective for ZIKV-infected mice (52).

Both ICompromised and ICompetent mice showed increased expression of placental *Tnf* mRNA, which has been demonstrated to be directly related to placental damage, abortion and premature birth (53–56). In addition, an increased *Tnf* response is related to impaired placental hormone production and trophoblastic invasion and increased apoptosis in pregnancy (57,58). Although we did not assess TNF-α protein levels in the placenta and maternal blood, this response could be implicated in the overall damage detected.

Although differences in placental weight were not observed, ZIKV infection mid-pregnancy had a profound effect on placental cellular turnover, dependent on titer, strain and/or placental compartment. The Lz is responsible for fetal and maternal nutrient, gas and waste exchange, while the Jz provides structural support, nutrient storage and hormone synthesis (35). ZIKV induced a consistent increase in Lz proliferation in all groups. However, the Lz apoptotic rate was increased only in the high-ZIKV-ICompetent and ICompromised groups and decreased in low-ZIKV-ICompetent mice. The mechanisms underlying these differences are unknown but may be related to the distinct maternal and placental proinflammatory responses and/or to the direct effect of the virus on the placenta (59). Increased Lz apoptosis in the high-ICompetent-ZIKV group may be one of the mechanisms driving the lower fetal and fetal head weight detected in this group. In this context, changes in placental turnover can determine placental maturation and function and lead to fetal distress and developmental abnormalities (60). An increase in the Lz apoptotic ratio may signify damage to this placental layer, which is consistent with the fact that diverse pathological lesions associated with congenital disorders were described in placentae from women infected by ZIKV at different stages of pregnancy (61). Conversely, no proliferative changes were observed in the Jz in high-ZIKV and low-ZIKV ICompetent mice, while increased and decreased apoptotic rates were detected. It follows that the lack of Jz-Ki-67 induction may suggest that this layer is less capable of restoring proliferation in response to high-ZIKV challenge, and this may be related to decreased fetal growth.

Our placental ultrastructural analysis detected consistent differences across ZIKV-exposed groups. The Lz and Jz layers from both strains exhibited signs of ER stress, i.e., dilated ER cisterns or fragmented ER granular structures. These alterations may result from the accumulation of folded or poorly folded viral proteins in the ER lumen (42,62,63). The *Flaviridae* family uses the ER to replicate (64), and according to Offerdahl et al. (2017)(63), there is evidence of ZIKV interacting with this organelle, promoting an increased release of Ca^+2^ from the ER to the cell cytoplasm, thereby causing an increase in the production of reactive oxygen species (ROS) (65,66).

The mitochondrial ultrastructure in the Lz and Jz layers was severely impacted by ZIKV exposure. We found evidence of mitochondrial degeneration, i.e., mitochondrial membrane rupture, absence of mitochondrial ridges and a less electron-dense mitochondrial matrix, in all the treated groups. Placental mitochondrial dysfunction is associated with IUGR (67,68) and may be related, at least in part, to the lower fetal weight observed in high-ZIKV-ICompetent fetuses along with the altered placental apoptotic and proliferative patterns. Furthermore, mitochondrial dysfunction together with ER stress is likely to modify the placental ROS balance and generate local oxidative stress (69), which is associated with impaired fetal development (70). Of importance, associations between mitochondrial disruption, ER stress and placental cell senescence have been reported. Senescence is characterized as an irreversible interruption of the cell cycle and acquisition of a senescence-associated secretory phenotype (SASP) that promotes the release of cytokines, such as IL-1, IL-6, IL-8 and proinflammatory proteases (70). Therefore, the increased expression of IL-6 detected in the placentas of ICompromised mice suggests a SASP profile, which may be related to changes in the ER and mitochondrial ultrastructure, accompanied by important changes in apoptosis and cell proliferation. The interactions between mitochondria and the ER are critical for homeostasis and cell signaling (71). In conjunction with the ER, mitochondria can regulate cell death mediators in response to hypoxia and inflammation (72). The increase in apoptosis observed in the high-titer ICompetent groups and the low-titer ICompromised group may be related to the mitochondrial damage and ER stress observed. In fact, we observed an important decrease in microvillus abundance in sinusoidal giant trophoblast cells. Previously, we observed a decrease in microvillus density in the Lz of pregnancies exposed to malaria in pregnancy (MiP) (36). Together, our data show that different gestational infective stimuli (MiP and ZIKV) are capable of damaging placental microvillus abundance and impairing proper fetal-maternal exchange function and fetal growth/survival.

Next, to investigate whether maternal ZIKV exposure may influence fetal protection, we evaluated the placental localization and expression (semiquantitative) of the ABC efflux transporter systems P-gp, Bcrp, Abca1 and Abcg1, which are highly enriched in labyrinthine microvilli and in human syncytiotrophoblasts. These efflux transporters exchange drugs, environmental toxins, cytotoxic oxysterols and lipids within the maternal-fetal interface (26). We found a consistent decrease in labyrinthine P-gp expression in all ZIKV-exposed groups, demonstrating that ZIKV infection during pregnancy has the potential to increase fetal exposure to P-gp substrates, such as synthetic glucocorticoids, antibiotics, antiretrovirals, antifungals, stomach-protective drugs, and nonsteroidal anti-inflammatory drugs (26). Furthermore, Jz-P-gp was decreased in ICompromised placentae. Although little is known about the function of ABC transporters in the Jz, our data highlight the need for further studies investigating the biological importance of ABC transporters in the placental endocrine and structural zones of the rodent hemochorial placenta under normal and infective conditions.

ZIKV impaired Lz Bcrp and Abca1 expression in ICompetent (high) and ICompromised (low) mice. However, no effects were observed in ICompetent animals at a low ZIKV titer or in Abcg1 in any experimental setting. Thus, ZIKV also likely increases fetal accumulation of Bcrp substrates (antibiotics, antiretrovirals, sulfonylureas, folate, mercuric species, estrogenic mycotoxins, carcinogens and phototoxic compounds, among others) and disrupts placental lipid homeostasis (lipids, cholesterol, and cytotoxic oxysterols) by reducing placental Abca1 expression (26,73–76). We can speculate that the increased fetal accumulation of the P-gp, Bcrp and Abca1 substrates during ZIKV infection may contribute to the establishment of congenital Zika syndrome, although additional studies are clearly required to answer this important question. The present data are in agreement with previous publications showing that bacterial, viral and protozoan inflammation alters the expression and/or function of P-gp, Bcrp and Abca1 in biological barriers, such as the placenta, yolk sac and blood-brain barriers (26,27,36,64,77–79).

## 5. Conclusion

Our data show that gestational ZIKV impacts the fetal phenotype independently of term fetal viremia. Abnormal placental cell turnover, ultrastructure and transporter expression may result from specific proinflammatory responses that depend on the ZIKV infective load and maternal immune status. Fetal accumulation of drugs, environmental toxins and lipids within the fetal compartment may potentially be increased in ZIKV-infected pregnancies due to altered levels of key ABC transporters.

## Author contributions

CBVA, FFB, EB, LBA and TMOC conceived and designed the experiments. CBVA, VRSM, SVAC, HRG, RPCS and VMON performed the experiments. CBVA, SVAC, FFB, EB, SGM, LBA and TMOC analyzed the data. CBVA, VRSM, EB, LBA and TMOC wrote the paper and edited the manuscript. All authors contributed to the article and approved the submitted version.

## Acknowledgments

We would like to thank Alan Moraes for supporting the acquisition of electron microscopy images and the electron microscopy laboratory at the UFF Biology Institute for allowing the use of the JEM1011 transmission electron microscope. We would also like to thank Mauro Jorge Castro Cabral for the use of the MAGPIX^®^ System equipment at the Paulo de Góes Institute of Microbiology/UFRJ; Ernesto T. Marques Jr. (Centro de Pesquisa Aggeu Magalhães, FIOCRUZ, PE) for providing ZIKV to the Institute of Microbiology Paulo de Góes, Federal University of Rio de Janeiro; and the technicians Juliana Gonçalves and Rakel Alves for their support during all the experiments.

